# Apical Constriction Reversal upon Mitotic Entry Underlies Different Morphogenetic Outcomes of Cell Division

**DOI:** 10.1101/862821

**Authors:** Clint S. Ko, Prateek Kalakuntla, Adam C. Martin

## Abstract

During development, coordinated cell shape changes and cell divisions sculpt tissues. While these individual cell behaviors have been extensively studied, how cell shape changes and cell divisions that occur concurrently in epithelia influence tissue shape is less understood. We addressed this question in two contexts of the early *Drosophila* embryo: premature cell division during mesoderm invagination, and native ectodermal cell divisions with ectopic activation of apical contractility. Using quantitative live-cell imaging, we demonstrated that mitotic entry reverses apical contractility by interfering with medioapical RhoA signaling. While premature mitotic entry inhibits mesoderm invagination, which relies on apical constriction, mitotic entry in an artificially contractile ectoderm induced ectopic tissue invaginations. Ectopic invaginations resulted from medioapical myosin loss in neighboring mitotic cells. This myosin loss enabled non-mitotic cells to apically constrict through mitotic cell stretching. Thus, the spatial pattern of mitotic entry can differentially regulate tissue shape through signal interference between apical contractility and mitosis.

## Introduction

Tissues grow in size and undergo complex morphogenetic movements to sculpt the embryo (LeGoff and Lecuit, 2015). Two major cell processes that contribute to morphogenesis are cell division and cell shape change. Often, these behaviors occur concurrently in the same tissue, leading to a complex interplay that can facilitate tissue-scale movements and shape changes (Etournay et al., 2015; Guirao et al., 2015; Li et al., 2014; Mao et al., 2013). For example, during the development of the *Drosophila* tracheal placode, cell division in the placode promotes fast cell internalization (Kondo and Hayashi, 2013). Cell divisions also drive cell rearrangements for proper gastrulation movements in the chick (Firmino et al., 2016) and promote tissue spreading during zebrafish epiboly (Campinho et al., 2013).

Apical constriction is a cell shape change that promotes tissue invagination (Leptin and Grunewald, 1990; Sawyer et al., 2010). During *Drosophila* gastrulation, the presumptive mesoderm cells on the ventral side of the embryo are internalized through coordinated apical constrictions to form the ventral furrow (Leptin and Grunewald, 1990; Sweeton et al., 1991). Apical contractility is activated by embryonic transcription factors Snail and Twist, which define mesoderm fate and also activate non-muscle myosin 2 (myosin) contractility through the small GTPase RhoA at the apical surface of cells (Costa et al., 1994; Dawes-Hoang et al., 2005; Kölsch et al., 2007; Young et al., 1991). In contrast to cases where cell divisions promote morphogenesis (Firmino et al., 2016; Kondo and Hayashi, 2013), premature mitotic entry during mesoderm invagination disrupts internalization (Großhans and Wieschaus, 2000; Mata et al., 2000; Seher and Leptin, 2000). Thus, cell division is actively repressed in the mesoderm. The *tribbles* (*trbl*) gene is one ventral-specific inhibitor of mitosis. In *trbl* mutants, cells in the prospective mesoderm prematurely divide, which disrupts mesoderm invagination (Großhans and Wieschaus, 2000; Mata et al., 2000; Seher and Leptin, 2000). This phenotype demonstrated the importance of coordinating cell shape change with cell cycle regulation, but it was not known how cell division disrupts mesoderm invagination. For example, without live-cell imaging it was unclear whether cell division prevents apical constriction from initiating and/or whether it interferes with apical constriction after it has started.

After 13 rounds of synchronous divisions in the early *Drosophila* embryo, the 14^th^ cycle of mitotic divisions occurs in a stereotypical pattern across the blastula, called mitotic domains, which correspond to regions of *string* (*stg*) expression (Edgar and Datar, 1996; Edgar and O’Farrell, 1989, 1990; Farrell and O’Farrell, 2014; Foe, 1989). String is the *Drosophila* homolog of Cdc25, a protein phosphatase that reverses inhibitory phosphorylation on cyclin-dependent kinase (Cdk1) (Gould et al., 1990; Russell and Nurse, 1986). Tribbles acts to degrade String protein in the mesoderm (Mata et al., 2000). While ventral fate-specific mitotic inhibition promotes mesoderm internalization, how the geometry and timing of mitotic entry influences cell and tissue shape change in other regions of the embryo is unknown.

Here, we determined how different spatial patterns of mitotic entry interact with apically constricting cells to affect tissue shape. In both native and artificially induced contractile epithelia, mitotic entry disrupts medioapical myosin activation and abrogates apical constriction. In the mesoderm, this disrupts tissue internalization. We showed that disruption of apical contractility is not due to loss of cell adhesion or apicobasal polarity but depends on mitotic entry. In contrast, ectopically contractile cells in the dorsal ectoderm situated between mitotic domains only apically constricted and invaginated when neighboring cells entered mitosis. In this context, internalization was associated with a force imbalance resulting from the loss of medioapical contractility in mitotic cells that neighbor contractile, non-mitotic cells. These results indicate that distinct morphogenetic outcomes result from different spatiotemporal patterns of mitotic entry and resulting changes in force generation.

## Results

### Premature mesodermal mitotic entry in *trbl* mutant embryos prevents or reverses anisotropic apical constriction

Previous studies used fixed embryos to study the *trbl* mutant phenotype so that it was not known how cell division disrupts mesoderm invagination. Therefore, to determine whether cell division prevents apical constriction from starting and/or impedes apical constriction after it has initiated, we imaged the apical surface of *trbl* mutant mesoderm cells in real time. We first verified the effectiveness of *trbl* RNA interference (RNAi) by imaging live embryos labeled for Histone::GFP (H2A::GFP) and membranes (Gap43::mCherry). Histone::GFP allowed us to visualize chromosome condensation, which marked mitotic entry. Consistent with previous work, *trbl* RNAi knockdown resulted in premature cell divisions in the mesoderm and a failure to form the ventral furrow (9/16 embryos) (Figure 1, A and B; Video 1) (Großhans and Wieschaus, 2000; Mata et al., 2000; Seher and Leptin, 2000). The timing of mitotic entry was variable in *trbl* RNAi embryos, which allowed us to determine the effects of mitotic entry when it happens either before or after apical constriction onset (Figure 1, B and C).

**Figure 1.**
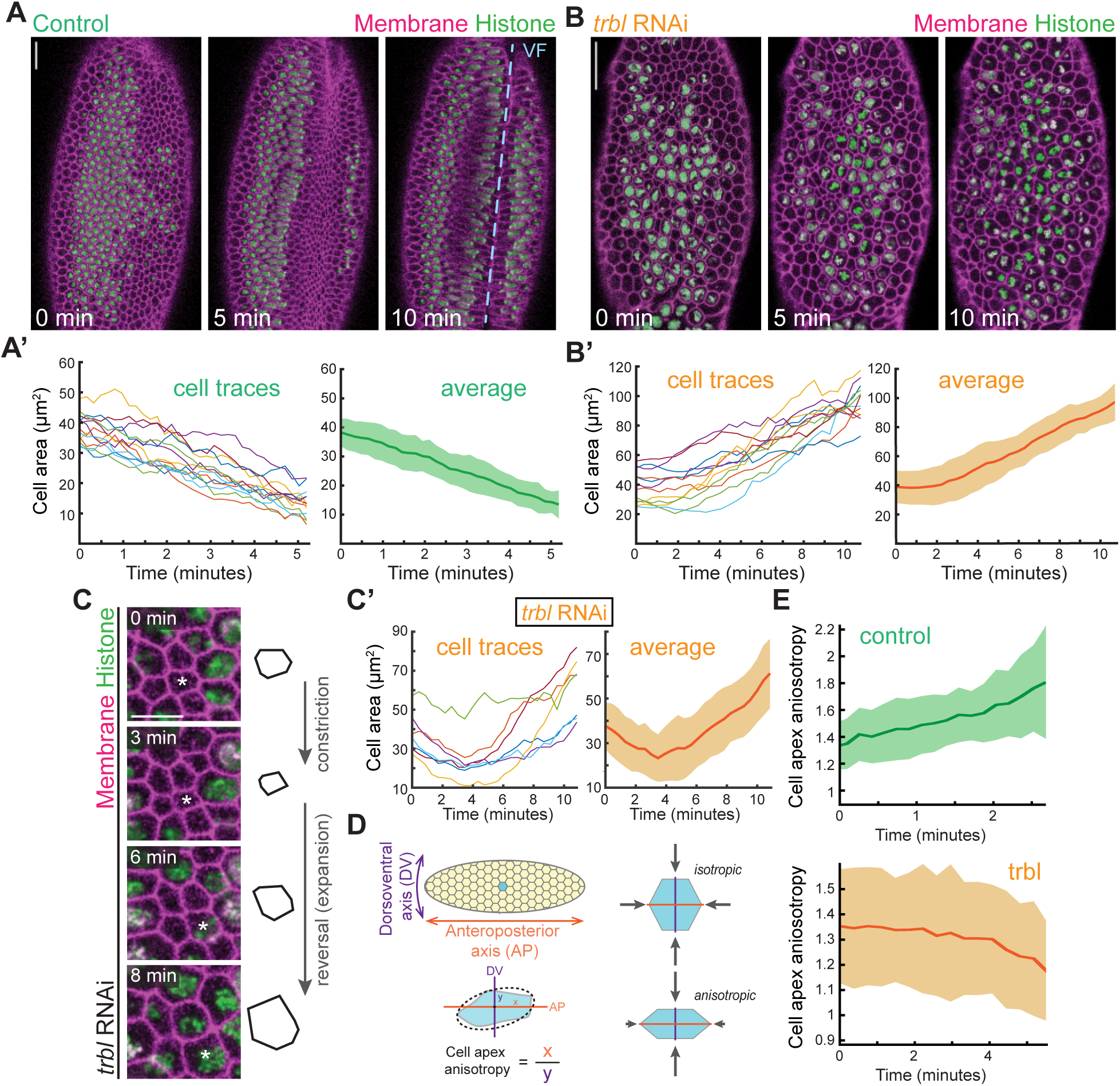
Premature mitotic entry in *trbl* mutant embryos reverses apical constrictions. (A-A’) During wild-type ventral furrow (VF, blue dashed line) formation, cells apically constrict. (A) Images are maximum intensity projections from a live embryo expressing H2A::GFP and Gap43::mCherry. (A’) Representative cells were segmented and their apical cell areas were tracked over time. The average trace of 12 cells with standard deviation is shown on the right. (B-B’) In *trbl* RNAi embryos, mesoderm cells prematurely divide and increase apical area. (B) Images are maximum intensity projections from a live embryo expressing H2A::GFP and Gap43::mCherry injected with *trbl* dsRNA. (B’) Representative cells were segmented and their apical cell areas were tracked over time. The average trace of 12 cells with standard deviation is shown on the right. (C) Individual cells in *trbl* embryos can initiate constriction and reverse their constricted shape upon mitotic entry. Images are maximum intensity projections from a live embryo expressing H2A::GFP and Gap43::mCherry injected with *trbl* dsRNA. An outline of the cell marked by the asterisk in the images is shown on the right. (C’) Quantification of changes in cell area for cells that initiate but reverse constriction after mitotic entry. Individual traces of 9 cells over 2 representative embryos injected with *trbl* dsRNA and the average with standard deviation are plotted. (D) Cartoon diagram depicting isotropic and anisotropic constrictions. Cell apex anisotropy is calculated as the cell length along the anteroposterior axis (AP, x) over the dorsoventral axis (DV, y). (E) Dividing cells in *trbl* RNAi embryos become more isotropic. Quantification of cell apex anisotropy over time in control and *trbl* RNAi embryos (after apical constriction has initiated). Scale bars, 20 μm (A and B), 10 μm (C).

To quantify the effect of mitotic entry, we segmented representative embryos from these data sets. Normally, apical constriction of the mesoderm is associated with tissue invagination (Figure 1A’) (Costa et al., 1994; Leptin and Grunewald, 1990; Sweeton et al., 1991). In contrast to control embryos, mesoderm cells in *trbl* RNAi embryos increased apical cell area as a consequence of mitotic rounding, a common phenomenon observed in non-constricting epithelial cells (Champion et al., 2016; Luxenburg et al., 2011; Reinsch and Karsenti, 1994; Rosa et al., 2015), which disrupted invagination (Figure 1, B and B’; Video 1). Because the timing of premature mitotic entry was variable such that not all cells synchronously divided, in several cases we found individual cells and embryos that had initiated apical constriction that then reversed their constricted shape and underwent apical expansion (Figure 1, C and C’). Thus, mitotic mesoderm cells do not sustain apical constriction.

An important feature of mesoderm cell apical constriction is that it is anisotropic, with greater constriction along the dorsoventral axis, which is important for inward tissue curvature and invagination (Figure 1D) (Chanet et al., 2017; Heer et al., 2017). This is reflected in the gradual increase of cell apex anisotropy (Figure 1D, anisotropy > 1) in control embryos after cells have initiated apical constriction (Figure 1E). However, in *trbl* embryos, after initial anisotropic constrictions, cell anisotropy decreased and approached a value of 1 due to mitotic rounding (Figure 1E). These results suggested that premature mitotic entry in the mesoderm can either prevent apical constriction from initiating or reverse apical constriction that has already started, depending on the timing of mitotic entry.

### Mitotic entry disrupts medioapical myosin activation

Apical constriction and mitotic rounding are dependent on actomyosin-based contractility (Dawes-Hoang et al., 2005; Kunda et al., 2008; Maddox and Burridge, 2003; Matthews et al., 2012; Rosa et al., 2015; Young et al., 1991). In the mesoderm, this involves an organized contractile machine with myosin enriched near the middle of the apical domain, the medioapical cortex (Mason et al., 2013; Coravos and Martin, 2016). To determine how premature mitotic entry in *trbl* mutants affected medioapical myosin, we imaged live embryos that were trans-heterozygous for a deficiency (Df(3L)ri79c) and a P-element insertion (EP(3)3519) that disrupt the *trbl* gene, which has previously been shown to exhibit the *trbl* mutant phenotype (Großhans and Wieschaus, 2000; Seher and Leptin, 2000). In contrast to wild-type or heterozygote embryos, which accumulate and sustain medioapical myosin, medioapical myosin failed to accumulate in Df/EP3519 embryos, with myosin instead localizing to junctional interfaces (Figure 2, A and B; Video 2). Despite initiating myosin accumulation, medioapical myosin was not sustained in ventral cells that entered mitosis (Figure 2, A and B; Video 2). We obtained a similar absence of medioapical myosin accumulation when we overexpressed *string* (Cdc25) in the early embryo (Figure S1A), which phenocopies *trbl* embryos (Großhans and Wieschaus, 2000; Seher and Leptin, 2000). In both *trbl* mutant and *string* overexpression embryos, medioapical myosin re-accumulated in ventral cells after completion of mitosis (Figure S1; Video 2). Thus, medioapical myosin activation is disrupted in ventral cells that prematurely enter mitosis, consistent with the observed increases in apical cell area (Figure 1, B and B’).

**Figure 2.**
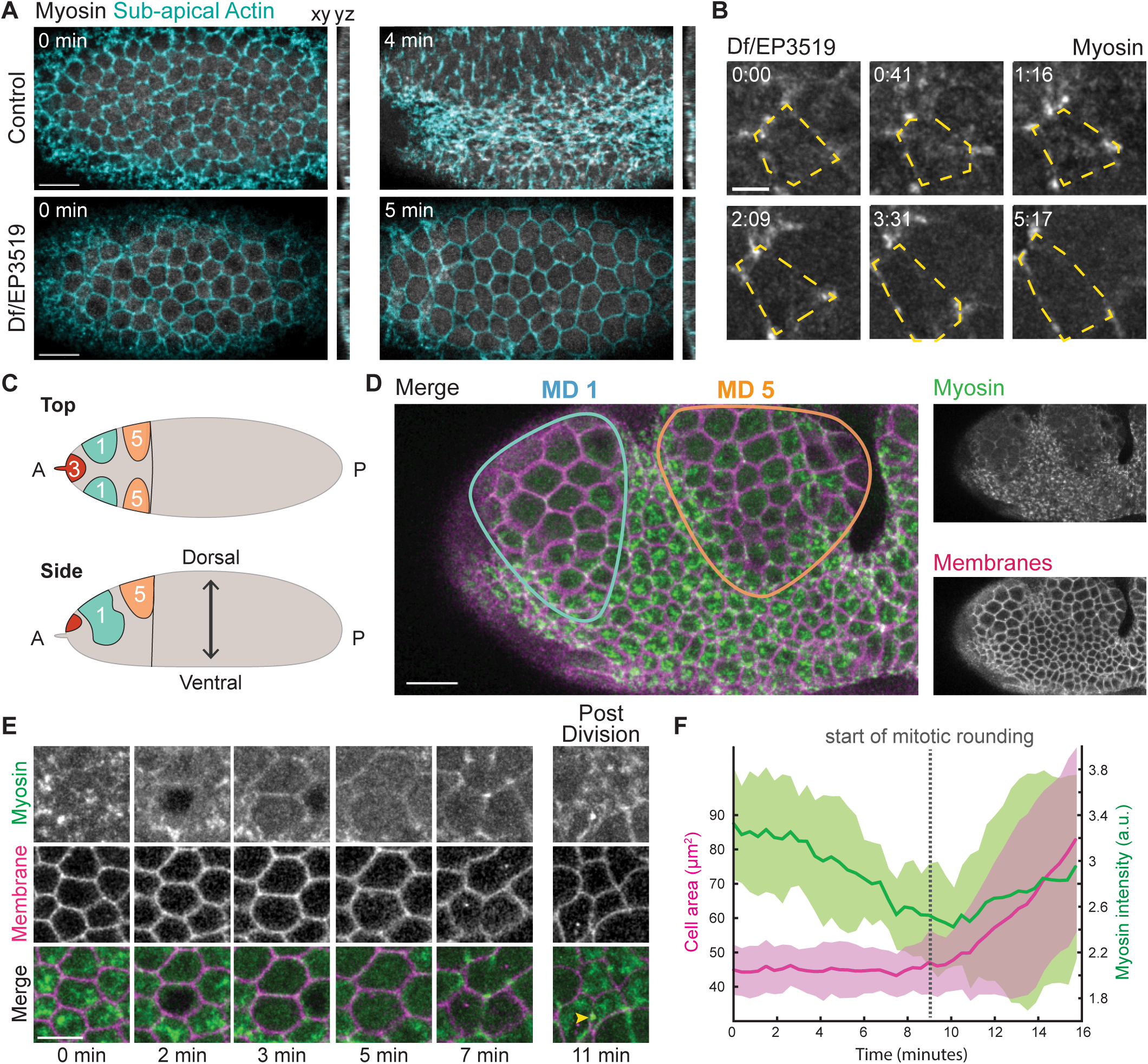
Apical myosin activation is disrupted upon mitotic entry. (A) Apical myosin is disrupted in cells that prematurely divide in *trbl* embryos. Images are maximum intensity projections from a live trans-heterozygous embryo (Df/EP3519) expressing Myo::CH (Sqh::CH) and Utr::GFP (Actin). Sub-apical sections of the Utr::GFP channel were used to mark cell outlines. Control embryos are heterozygotes with a wild-type copy of *trbl*. (B) Montage of a Df/EP3519 embryo expressing Myo::GFP, which shows apical myosin dissipates as cells round. The outline of one cell is highlighted by yellow dashed lines. The cell outline was determined from the sub-apical myosin signal. (C) Cartoon diagram showing mitotic domains (MD) 1,3, and 5 (blue, red, and orange, respectively). (D) Apical myosin is lost in mitotic domain cells in the ectoderm. Images are maximum intensity projections from a live embryo with ectopic *fog* in the ectoderm expressing Myo::GFP and Gap43::mCherry. (E) Montage of *fog-* overexpressing embryo with Myo::GFP and Gap43::mCherrry. Apical myosin reaccumulates in both daughter cells after mitosis completes. Midbody is marked by the yellow arrowhead. (F) Quantification of mean cell area (lilac) and mean myosin intensity (green) with standard deviations for a representative *fog-*overexpressing embryo (n = 10 cells). Scale bars, 15 μm (A and D), 10 μm (E), 5 μm (B).

To determine whether loss of medioapical myosin was a general feature of dividing, contractile epithelial cells, we took advantage of the stereotyped cell divisions in the early mitotic domains that occur on the dorsal side of the head (Foe, 1989), particularly focusing on mitotic domains 1 and 5 (Figure 2C). We artificially increased ectoderm apical contractility by ectopically expressing *folded gastrulation* (*fog*), a ligand for a G-protein-coupled receptor (GPCR) that is expressed in the mesoderm and functions upstream of apical myosin activation (Costa et al., 1994; Dawes-Hoang et al., 2005; Manning et al., 2013; Sweeton et al., 1991). However, a GPCR for Fog is also present in the ectoderm and ectopic *fog* expression in this tissue leads to apical myosin accumulation (Dawes-Hoang et al., 2005; Kerridge et al., 2016). This allowed us to upregulate apical myosin levels consistently prior to mitotic entry and to compare apical myosin levels in mitotic and non-mitotic cells in the same tissue without interfering with the normal developmental progression of cell divisions in the embryo. Similar to *trbl* mutant embryos, Fog-induced medioapical myosin decreased in mitotic cells (Figure 2, D–F; Video 3). As medioapical myosin spots dissipated, myosin localization became isotropically localized around the cell cortex, a feature of mitotic rounding (Figure 2, D and E) (Maddox and Burridge, 2003; Matthews et al., 2012; Ramanathan et al., 2015; Rosa et al., 2015; Stewart et al., 2010). The medioapical myosin meshwork returned in both daughter cells after mitotic exit and cytokinesis (Figure 2E). These results suggested that mitotic entry temporarily overrides cell type-specific signaling in both mesoderm and ectoderm that promotes apical contractility.

### Medioapical myosin disruption is not due to loss of cell adhesion or apicobasal polarity

Because cells round up upon disruption of adherens junctions (Martin et al., 2010), it was possible that mitotic entry disrupted intercellular adhesion. However, the disruption of medioapical myosin preceded the apical cell area expansion (i.e., rounding) (Figure 2F), suggesting the apical myosin loss is not caused by disrupted adhesion. To test whether changes in myosin regulation were dependent on changes in cell shape or adhesion during cell division, we disrupted cell adhesion with a maternal and zygotic loss-of-function mutant in the *Drosophila* β-catenin gene (*armadillo, arm*) and analyzed mitotic progression. The *arm* mutant disrupts the mechanical integrity of tissues, causing constitutively round cells that do not invaginate (Cox et al., 1996; Dawes-Hoang et al., 2005). However, even when cell adhesion was lost and individual cells became rounded, apical contractility was sustained (Figure 3, A and B) (Dawes-Hoang et al., 2005; Martin et al., 2010).

**Figure 3.**
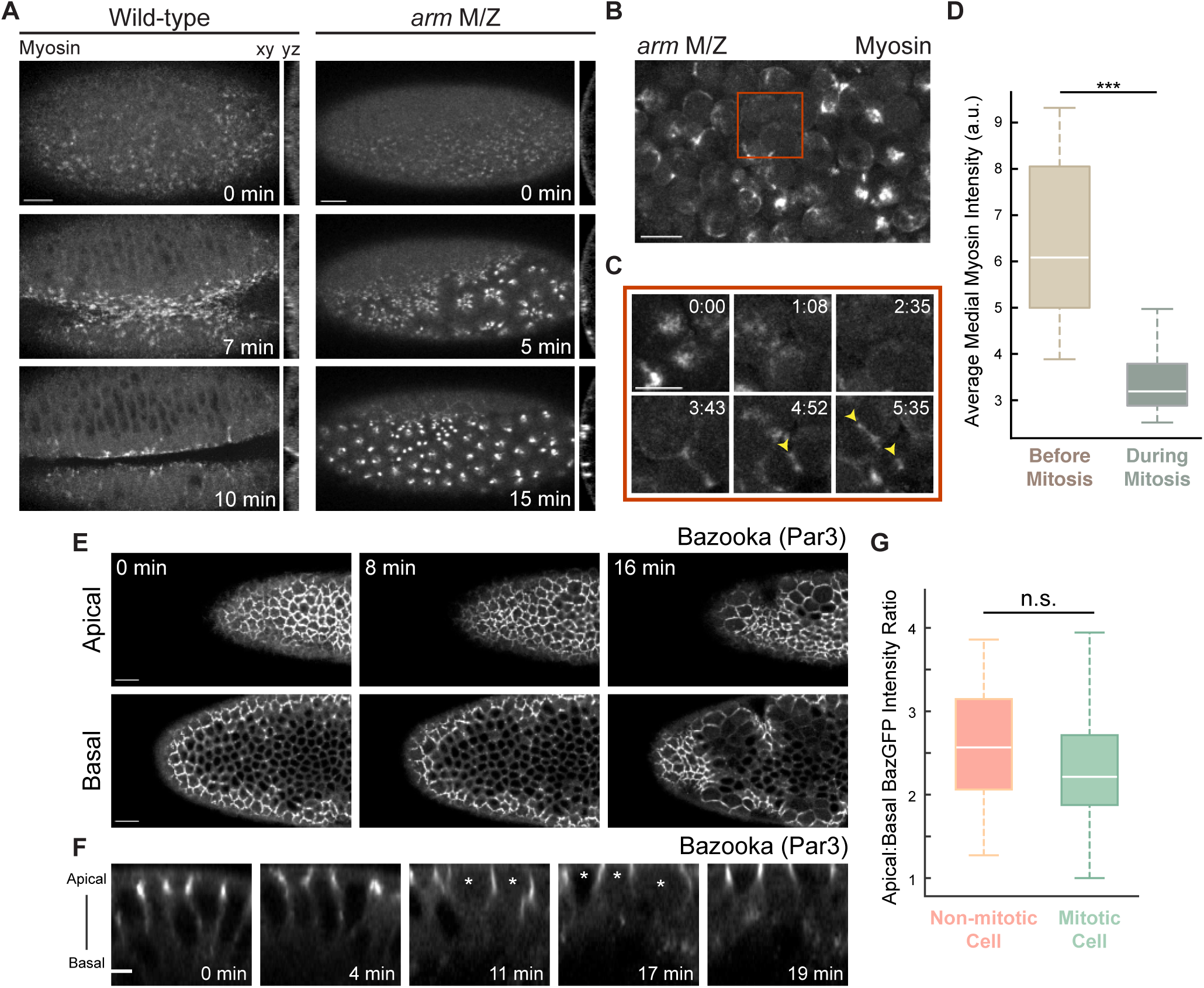
Apical contractility loss is not due to disrupted adhesion or apical-basal polarity. (A) In *arm* mutants, cells become mechanically uncoupled and the supracellular myosin meshwork fragmented. Images are maximum intensity projections from control (wild-type) and maternal and zygotic *arm* mutants expressing Myo::GFP. Cross-section views are to the right of each *en face* view. (B) Apical contractility is sustained in cells with rounded morphology in *arm* mutants. Image is a maximum intensity projection from a live maternal and zygotic *arm* mutant expressing Myo::GFP. (C) Apical myosin is lost during mitosis in rounded *arm* mutant cells. Images are maximum intensity projections from a maternal and zygotic *arm* mutant expressing Myo::GFP (magnified images from red box in B, starting at a later time point). Cytokinetic furrows are highlighted by yellow arrowheads. (D) Quantification of average medial myosin intensity in *arm* mutant cells. Myosin intensity was measured before mitosis by selecting cells just prior to nuclear envelope breakdown and compared to myosin intensities in cells just prior to cytokinesis (n = 20 cells; ***, P < .0001, unpaired t test). Bottom and top edges of the boxplot are 25^th^ and 75^th^ percentiles, with median marked by the white line. Whiskers extend to the most extreme data points. (E) Baz polarity is unaffected during mitosis. Images are apical (top) and basal (∼8 μm below apical slice; bottom) *en face* views of embryos with ectopic *fog* expressing GFP-tagged Baz. Cross-section views of mitotic cells are shown in (F). Mitotic cells are marked with white asterisks. (G) Baz is apically polarized in both mitotic and non-mitotic cells. Quantification of the ratio of maximum pixel intensity values of Baz::GFP in the apical to basolateral domain (n = 30 cells each across 3 embryos; unpaired t test). Bottom and top edges of the boxplot are 25^th^ and 75^th^ percentiles, with median marked by the white line. Whiskers extend to the most extreme data points. Scale bars, 15 μm (A, B, and E), 10 μm (C), 5 μm (F).

During gastrulation, cell division normally proceeds in mesoderm cells after they have internalized (Foe, 1989). However, because *arm* mutants block invagination, we could examine the consequence of mitotic entry on non-adherent cells at the embryo surface. In *arm* mutants, apical myosin spots disappeared only when the mesoderm cells entered mitosis even though cells had maintained a rounded morphology prior to mitoses (Figure 3, C and D) (Foe, 1989). Thus, the switch in myosin regulation is independent of changes in cell shape and adhesion, suggesting that mitotic entry disrupts other processes that are required for apical contractility.

Alternatively, we hypothesized that apical contractility defects could be due to a loss of apicobasal polarity. To test this, we determined if mitotic entry of ectodermal cells in embryos with ectopic *fog* expression affected the apical-basal polarity of Bazooka (Baz, Par3), a component of the apical polarity complex that plays an important role in establishing and maintaining apicobasal polarity (Bilder et al., 2003; Harris and Peifer, 2004, 2007). In cells of embryos with ectopic *fog* expression, Baz was localized to apical junctions (Figure 3, E–G). However, in mitotic cells, polarized Baz localization was retained during mitotic rounding (Figure 3, F and G), suggesting that loss of medioapical myosin at the onset of mitotic rounding was also not due to a loss of apicobasal polarity.

### Mitotic entry in apically constricting cells changes RhoA regulation

To determine the basis for mitosis-dependent changes in myosin localization, we examined RhoA activity in *trbl* mutants and in the early mitotic domains of embryos with ectopic *fog* expression. Apical constriction and mitotic rounding involve RhoA activation downstream of the Rho guanine nucleotide exchange factors (GEFs), RhoGEF2 and Ect2/Pebble (Pbl), respectively (Barrett et al., 1997; Häcker and Perrimon, 1998; Kölsch et al., 2007; Maddox and Burridge, 2003; Matthews et al., 2012; Rosa et al., 2015; Yoshizaki et al., 2003). As a marker for RhoA activity, we first examined the localization of a GFP-tagged Rho-associated coiled-coil kinase (ROCK) (Simões et al., 2010), the RhoA effector that exhibits RhoA-dependent medioapical cortex localization during apical constriction (Mason et al., 2016). ROCK phosphorylates and activates myosin (Amano et al., 1996; Mason et al., 2013; Mizuno et al., 1999; Royou et al., 2002). In Df/EP3519 *trbl* mutant embryos, medioapical ROCK localization either did not accumulate or was lost in mesoderm cells when they prematurely entered mitosis (Figure 4, A and B). Thus, mitotic entry disrupted medioapical ROCK localization associated with apical constriction, suggesting a disruption of medioapical RhoA activity.

**Figure 4.**
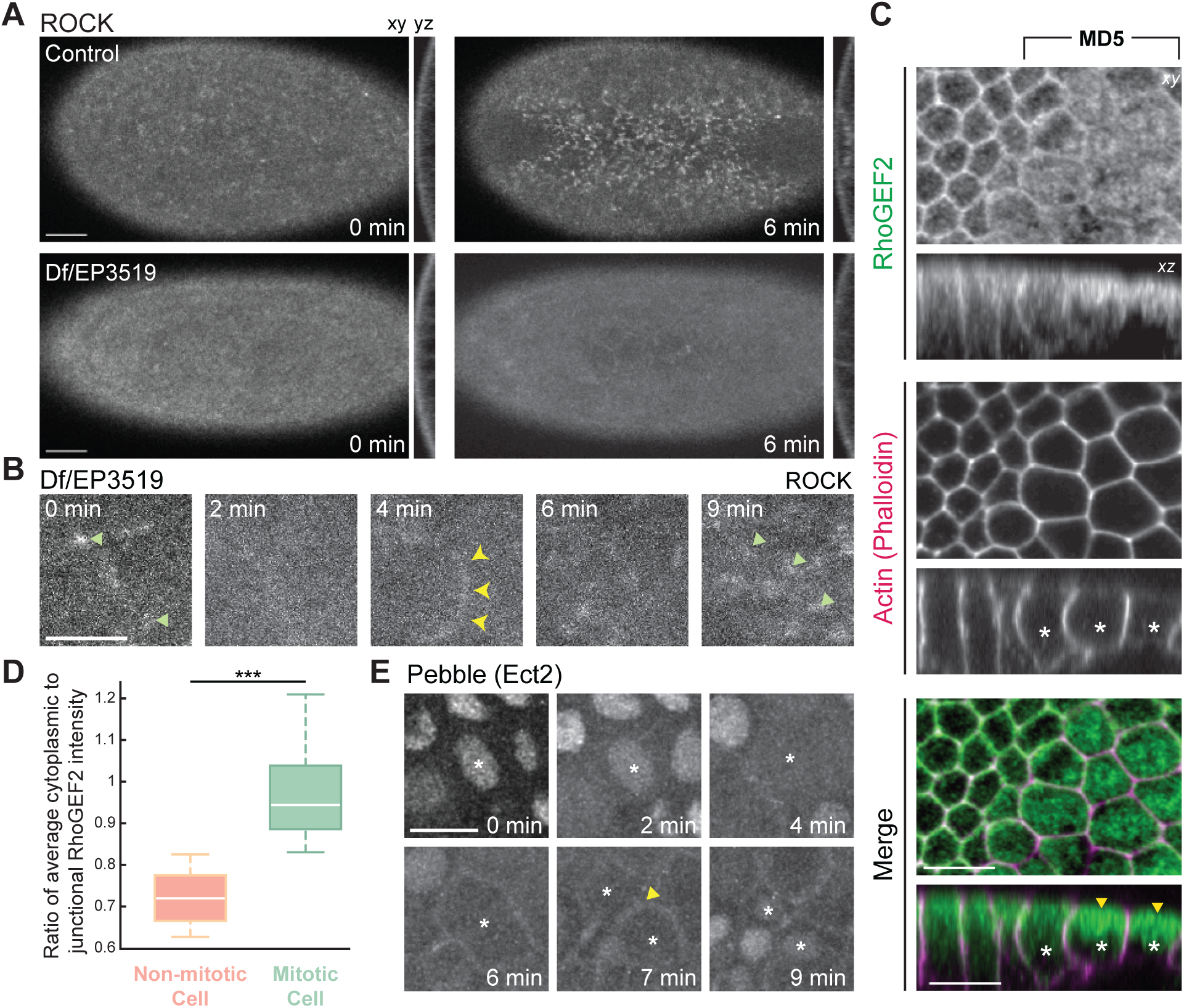
Different Rho GEFs exhibit distinct localization changes upon mitotic entry. (A) Medioapical ROCK localization is not sustained in *trbl* mutants. Images are maximum intensity projections from embryos that are either trans-heterozygous for the deficiency and P-element insertion (mutant) or not trans-heterozygous (control). Embryos are also expressing kinase dead rok(K116A)::GFP. (B) Apical ROCK foci (green arrows) disappear during mitosis. Montage from the *trbl* mutant embryo shown in (A). A cytokinetic ring is highlighted by the yellow arrowheads. (C) RhoGEF2 localization is less cortical and more cytoplasmic in mitotic cells. Images are single sub-apical slices of a fixed representative *fog* overexpressing embryo immunostained against GFP-tagged RhoGEF2 and phalloidin to visualize F-actin. Asterisks mark mitotic cells in the cross-section images (bottom) with cytoplasmic enrichment of RhoGEF2 highlighted by yellow arrowheads. (D). RhoGEF2 becomes enriched in the cytoplasm in mitotic cells. Quantification of the ratio of average cytoplasmic to junctional RhoGEF2 intensity in mitotic or non-mitotic cells (n = 20 cells across 4 embryos; ***, P < .0001, unpaired t test). Bottom and top edges of the boxplot are 25^th^ and 75^th^ percentiles, with median marked by the white line. Whiskers extend to the most extreme data points. (E) Pebble/Ect2 localizes to the cortex after mitotic entry in mitotic domain 3. Images are maximum intensity projections from a live embryo expressing Pbl::GFP under a myosin promoter. One mitotic cell from a mitotic domain and its daughter cells are marked by the asterisks. The site of cytokinetic furrow formation is marked by the yellow arrowhead. Scale bars, 15 μm (A), 10 μm (B - E).

To determine how RhoA activity was disrupted, we investigated the localization of RhoGEFs that are associated with either apical constriction or mitotic rounding. We imaged mitotic domains in embryos ectopically expressing *fog*, due to technical challenges with combining GFP-tagged RhoGEFs with the *trbl* mutants. First, we fixed embryos with ectopic *fog* expression that also expressed GFP-tagged RhoGEF2 under an endogenous promoter and immunostained with an anti-GFP antibody. Immunofluorescence of fixed embryos gave us the clearest signal to visualize RhoGEF2 in mitotic cells because the autofluorescence of the vitelline membrane could be removed. Consistent with previous work in mesoderm cells, non-mitotic ectoderm cells ectopically expressing *fog* exhibited apically enriched, junctional RhoGEF2 (Figure 4C) (Kölsch et al., 2007; Mason et al., 2016). In contrast, there was a clear reduction of apico-junctional RhoGEF2 and an associated increase in cytoplasmic signal in mitotic cells (Figure 4C, yellow arrowheads; Figure 4D). In conjunction with the observed changes in RhoGEF2 localization, Ect2/Pbl relocalized from the nucleus to the cortex in mitotic domain cells and became enriched at the spindle midzone during cytokinesis, similar to what has been described for other non-apically constricting cells (Figure 4E) (Matthews et al., 2012; Rosa et al., 2015). These results suggested that Ect2/Pbl-mediated cortical contractility is distinct from medioapical contractility mediated by RhoGEF2 (Kölsch et al., 2007) and that changes in RhoGEF2 localization underlie the disruption of medioapical myosin activation.

### Mitotic entry flanking contractile tissue promotes invagination via downregulation of opposing force

While premature mitotic entry in the mesoderm inhibited invagination, we discovered that cell divisions in the dorsal head of embryos with ectopic *fog* expression promoted ectopic tissue invaginations (Figure 5, A and B; Video 3). Normally, mitotic domains do not result in furrow formation (Foe 1989), as shown in control embryos lacking ectopic *fog* expression (Figure S2). In contrast, when *fog* was ectopically expressed in embryos, ectopic furrows formed between mitotic domains in regions where cells maintained apical contractility (Figure 5, A and B; Video 3).

**Figure 5.**
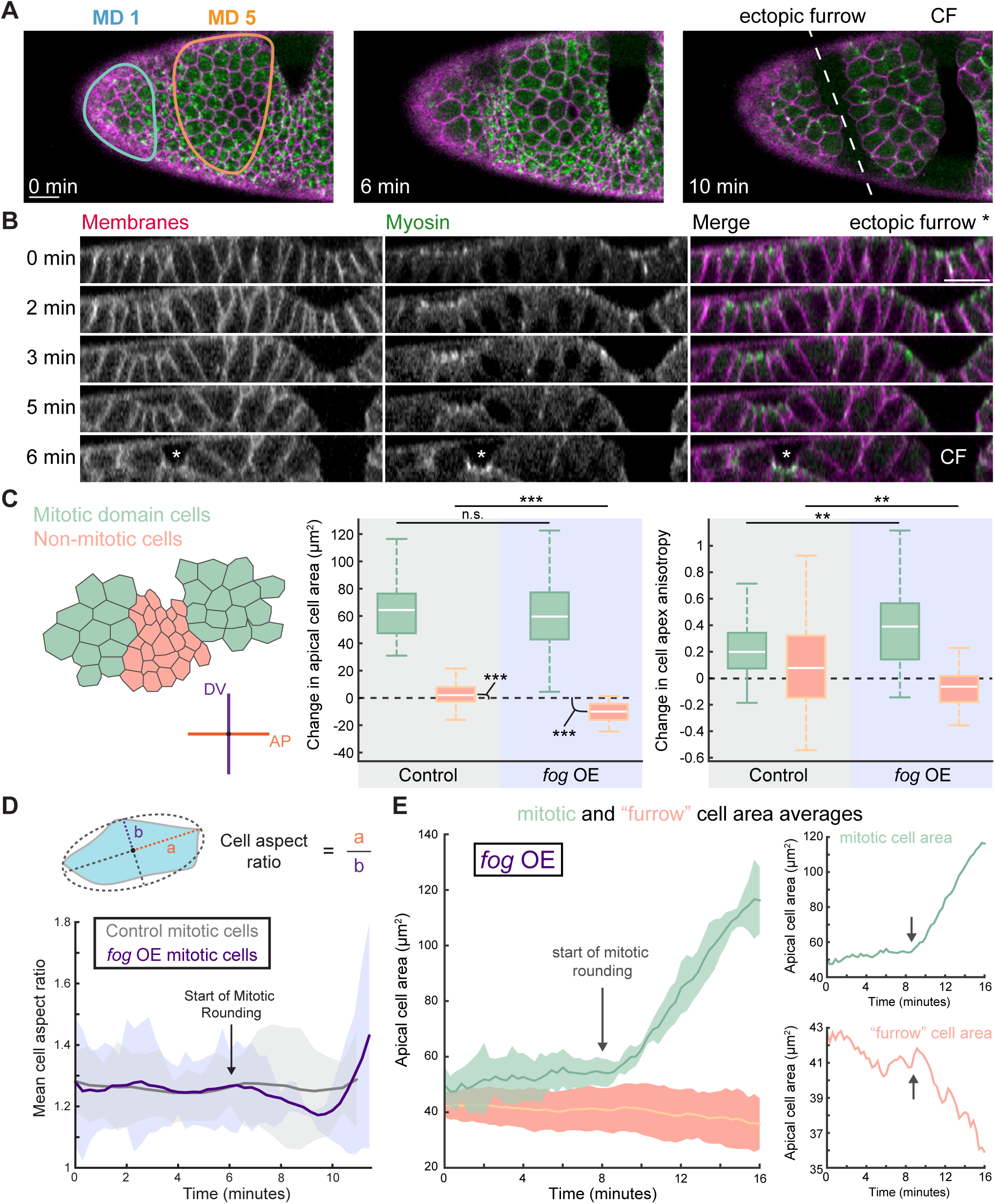
Ectopic furrows form between mitotic domains in embryos with ectopic *fog* expression. (A) Non-mitotic, contractile cells between mitotic domains invaginate during gastrulation. Images are maximum intensity projections from a live *fog* overexpressing embryo expressing Myo::GFP and Gap43::mCherry. The ectopic furrow is shown by a white dashed line. The invagination posterior to mitotic domain (MD) 5 is the cephalic furrow (CF). (B) Cross-section views of local tissue invaginations from the embryo in (A). Images from control embryo not ectopically expressing *fog* are in Figure S2. (C) Quantification of apical cell area and cell apex anisotropies in non-mitotic (salmon in cartoon) and mitotic (mint green in cartoon) cells. Final cell area was evaluated just before mitotic cells formed cytokinetic furrows. The time elapsed between initial and final cell area was between 5 and 20 minutes. Across 6 representative *fog* overexpressing embryos, 26 furrow cells and 33 mitotic domain cells were analyzed. Across 5 representative control embryos (Rhodopsin 3 shRNA line), 133 non-mitotic cells and 66 mitotic domain cells were analyzed (***, P < .0001; **, P < .01, unpaired *t* test). For changes in cell area, significance from 0 was determined with a one sample t test. Bottom and top edges of the boxplot are 25^th^ and 75^th^ percentiles, with median marked by the white line. Whiskers extend to the most extreme data points. (D) Cell aspect ratio increases more in embryos with ectopic *fog* expression. Quantification of mean change in cell aspect ratio with standard deviations between a representative control (Rhodopsin 3 shRNA line) and ectopic *fog* expression embryo. Cell aspect ratio is calculated as the distance from the centroid of a fitted ellipse to the ellipse edge along the major axis (a) over the distance along the minor axis (b). For ectopic *fog* expression embryos, 6 cells were quantified, and 7 cells were quantified for control embryos. Aspect ratio was measured up to the start of cytokinesis. (E) Apical constriction of non-mitotic cells initiate when neighboring mitotic domain cells enter mitosis. Quantification of apical cell area in a representative *fog* overexpressing embryo. Individual cell traces as well as averages with standard deviation are shown for mitotic domain cells (mint; n = 6 cells) and non-mitotic domain cells (salmon; n = 28 cells). The initiation of mitotic rounding is marked by the arrow. Scale bars, 15 μm.

To determine how furrows formed between mitotic domains, we analyzed the apical area of non-mitotic cells that formed the ectopic furrow. In control embryos, non-mitotic cells situated between mitotic domains did not exhibit a net decrease in apical area, presumably because these cells did not generate contractile force (Figure 5C). In contrast, cells between mitotic domains in embryos ectopically expressing *fog* underwent apical constriction (Figure 5, B and C, compare salmon boxes, left graph). Importantly, invagination onset was triggered when cells in the mitotic domains entered mitosis (Figure 5, A and B; Video 3), and these invaginations occurred before the completion of cytokinesis, suggesting that mitotic entry and not increased cell number promoted invagination. Thus, within a uniformly contractile tissue, domains of cells that enter mitosis can promote constriction and invagination of neighboring cells.

Because furrowing only occurred when the ectoderm was contractile, we tested how mitotic domains promote apical constriction in neighboring cells. One hypothesis is that furrowing could be due to isotropic pushing forces generated by mitotic rounding (Kondo and Hayashi, 2013). Alternatively, because mitotic entry reduces medioapical contractility, mitotic entry could downregulate force that opposes constriction and allow neighboring cells to change shape. Cell expansion or relaxation is important for morphogenesis in other contexts, often compensating for changes in neighboring tissue regions (Gutzman and Sive, 2010; Perez-Mockus et al., 2017; West et al., 2017). If the latter case is true, one prediction is that mitotic cells would stretch towards the ectopic furrow because of pulling forces from adjacent, contractile cells. Consistent with both hypotheses, the apical areas of mitotic cells increased to the same extent regardless of *fog* expression (Figure 5C, compare mint green boxes, left graph). However, mitotic domain cells in embryos with ectopic *fog* expression were more elongated and stretched towards the ectopic furrow with a greater increase in cell apex anisotropy than control embryos (Figure 5C, compare mint green boxes, right graph), suggesting that the intervening non-mitotic cells that apically constrict pull and stretch mitotic cells. In addition, we measured the cell aspect ratio (major/minor axis of fitted ellipse) and compared control and ectopic *fog* embryos. In both cases, mitotic cells exhibited an initial decrease in cell aspect ratio (due to cell rounding) but increased in aspect ratio prior to cytokinesis, likely during anaphase (Figure 5D) (Ramkumar and Baum, 2016). However, we found that mitotic cells from embryos with ectopic *fog expression* exhibited a greater change in cell aspect ratio (i.e., more elongation) (Figure 5D; Figure S3), suggesting that the higher anisotropy of dividing cells in tissues with ectopic contractility cannot be fully explained by normal anaphase elongation. Furthermore, the apical area of ectopic furrow cells only reduced after neighboring cells entered mitosis (Figure 5E), lending additional support for the idea that mitotic cell rounding and then elongation relative to neighboring non-mitotic cells, creates a force imbalance that allows neighboring cells to apically constrict and invaginate. These results indicated that the reversal of medioapical contractility and apical expansion that occurs during mitotic entry promotes tissue invagination when mitotic entry occurs adjacent to contractile cells (Figure 6).

**Figure 6.**
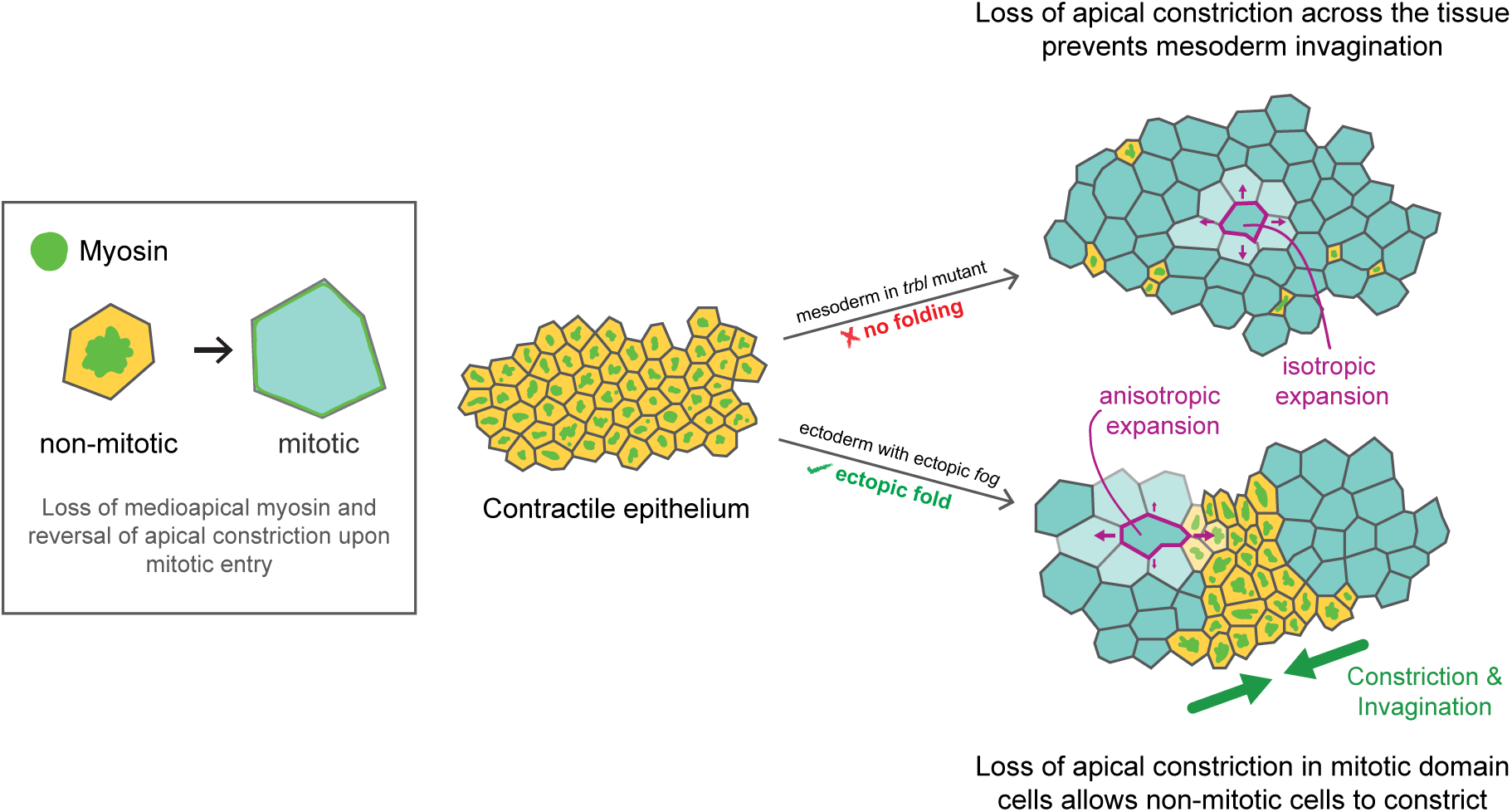
Different patterns of mitotic entry result in distinct morphogenetic outcomes. Cartoon diagram of a model contractile epithelium with different spatial patterns of mitotic entry. Apically constricting cells (yellow) that enter mitosis (blue) lose medioapical myosin and reverse their constricted cell shape (summarized in box). In the *trbl* mutant (top), most of the cells in the contractile tissue enter mitosis, which disrupts tissue folding. Mitotic cells in the mesoderm expand isotropically (magenta arrows). In contrast, when mitotic cells are interspersed by non-mitotic cells that sustain apical contractility, such as in the dorsal head of embryos with ectopic *fog* expression (bottom), mitotic cells that lose medioapical myosin expand anisotropically (magenta arrows) as they are pulled towards constricting cells (green arrows).

## Discussion

Here, we investigated the impact of mitotic entry in two different contractile epithelia with opposing tissue shape outcomes. Cell cycle-regulated changes in the cell, in particular the formation of an isotropic actomyosin cortex during mitotic rounding, is commonly observed across epithelial cell types and has been well-characterized (Maddox and Burridge, 2003; Matthews et al., 2012; Ramanathan et al., 2015; Rosa et al., 2015; Sorce et al., 2015; Stewart et al., 2010). However, it was previously unknown how mitotic entry would dynamically affect epithelial cells that are actively constricting. Through live imaging of apically constricting cells undergoing mitosis, we found that mitotic entry disrupts medioapical contractile signaling. In both the mesoderm of *trbl* mutants and the ectoderm with ectopic *fog* expression, medioapical myosin accumulation was reversed. We found that this change was followed by cell rounding and isotropic cortical myosin accumulation, which are specific to mitotic entry and not due to loss of cell adhesion. Indeed, previous work has demonstrated that mitotic progression in embryonic epithelial cells is only associated with local remodeling of cell adhesion at the site of cytokinesis, which allows epithelial integrity to be maintained (Founounou et al., 2013; Guillot and Lecuit, 2013; Herszterg et al., 2013; Higashi et al., 2016). The loss of medioapical myosin was not due to loss of cell adhesion or apicobasal polarity because mitotic downregulation of myosin still occurred in *arm* mutant germline clones and Baz localization remained apical throughout mitosis. Importantly, we also found that mitotic entry disrupts medioapical RhoA signaling and cortical RhoGEF2 localization, even though Ect2/Pbl becomes cortical, as previously reported (Matthews et al., 2012; Rosa et al., 2015).

We present a new paradigm for how cell divisions influence morphogenetic events: cell cycle-dependent changes in RhoA regulation can either inhibit or promote tissue shape change depending on differences in the spatiotemporal pattern of mitotic entry in the tissue. During mesoderm invagination, mitotic downregulation of medioapical contractility in the same cells that are needed to undergo apical constriction disrupted invagination (Großhans and Wieschaus, 2000; Leptin and Grunewald, 1990; Mata et al., 2000; Seher and Leptin, 2000; Sweeton et al., 1991). In contrast, mitotic downregulation of medioapical contractility in cells neighboring contractile cells promoted invagination. Here, we propose that medioapical myosin loss upon mitotic entry caused apical cortex relaxation relative to neighboring contractile cells. In support of this force imbalance model, mitotic cells elongate towards constricting cells prior to cytokinesis, leading to mitotic cell shape anisotropy that is higher than mitotic cells not neighboring contractile cells. In contrast, mitotic cells in the mesoderm of *trbl* mutants expanded their apical areas isotropically and remain isotropic through cytokinesis. Thus, cell cycle-mediated loss of medioapical myosin can be harnessed to provide local regions of tissue relaxation that can drive tissue folding.

Mitotic entry overrides or inhibits intracellular signaling that promotes the assembly of the medioapical contractile machine, remodeling the cytoskeleton in a way that leads to relaxation of the apical cortex. This creates a force imbalance where mitotic cells can become more compliant relative to their neighbors. This is similar to the idea that lateral ectoderm cells in the *Drosophila* embryo are less stiff, allowing the mesoderm to internalize (Perez-Mockus et al., 2017). Differences in epithelial tension also drive tissue folds in the *Drosophila* wing discs (Sui et al., 2018) and differential cell division and growth contribute to the positioning of these folds (Tozluoglu et al., 2019). In light of our results, it would be interesting to examine whether epithelial invagination in other contexts are bordered by cell divisions.

One potential molecular explanation for why medioapical myosin is lost during mitosis is that the two distinct cytoskeletal organizations that promote apical constriction or mitotic rounding compete for a limited pool of cytoskeletal components. Limited availability of actin monomers have been shown to play a role in how different actin network densities and sizes are regulated (Suarez and Kovar, 2016). For example, in fission yeast, inhibiting F-actin polymerization through the Arp2/3 complex results in an increase in formin-mediated F-actin assembly (Burke et al., 2014). However, given the apparent changes to RhoA signaling that occur in *fog* positive cells that enter mitosis, we favor a model in which signaling crosstalk or competition for upstream signals disrupts apical RhoA signaling (Agarwal and Zaidel-Bar, 2019; Jaffe and Hall, 2005).

To promote the assembly of medioapical actomyosin networks in the early *Drosophila* embryo, RhoGEF2 is the primary RhoA GEF (Barrett et al., 1997; Dawes-Hoang et al., 2005; De Las Bayonas et al., 2019; Fox and Peifer, 2007; Häcker and Perrimon, 1998; Kölsch et al., 2007). RhoGEF2 is thought to be particularly important for activating medioapical contractility (De Las Bayonas et al., 2019; Kerridge et al., 2016). To promote mitotic rounding, Ect2/Pebble is the primary RhoA activator (Matthews et al., 2012; Rosa et al., 2015). Our results indicate that these distinct Rho GEFs do not act additively. However, the precise nature by which RhoA activity is regulated downstream of RhoGEF2 and Ect2/Pebble in the same cell is still unclear. Activation of mitotic entry may affect RhoGEF2 localization because medioapical RhoGEF2 is influenced by microtubules and microtubule dynamics change in mitosis (De Las Bayonas et al., 2019; Rogers et al., 2004). However, disruption of microtubules does not prevent medioapical myosin activation (Ko et al., 2019). Mitotic entry may also affect signaling processes upstream of Rho GEF activation, such as the well-characterized case of GPCR signaling in *Drosophila* that activates different modes of contractility (Costa et al., 1994; Dawes-Hoang et al., 2005; Jha et al., 2018; Kerridge et al., 2016).

## Materials and Methods

### Fly stocks and genetics

Fly stocks and crosses used in this study are listed in Table S1. Crosses were maintained at 27 °C. In the F2 generation, non-balancer females and males were used to set up cages that were incubated at 25 °C. All other crosses and cages were maintained at 25 °C. To generate maternal and zygotic *arm* mutants expressing Myo::GFP, *arm*^034A01^ FRT101/FM7; sqh-GFP females were crossed to male *ovo^D^* FRT101/Y; hsFlp to obtain *arm*^034A01^ FRT101/ *ovo^D^* FRT101 females. These females were heat shocked at the larval stage at 37 °C for 2 hours over 3 to 4 days to induce mitotic recombination.

### Live and fixed imaging

For live imaging, embryos were dechorionated in 50% bleach, washed in water, and mounted onto a glass slide coated with glue (double-sided tape dissolved in heptane). Coverslips (No. 1.5) coated in glue were attached to the slide to use as spacers and a No. 1 coverslip was attached on top to create a chamber. Halocarbon 27 oil was used to fill the chamber. All imaging took place at room temperature (∼ 23 °C).

For fixed imaging, embryos with ectopic *fog* expression and control (Rhodopsin-3 shRNA line) embryos were dechorionated in bleach, washed in water, and fixed in 8% paraformaldehyde in 0.1 M phosphate buffer at pH 7.4 with 50% heptane for 30 min and manually devitellinized with a 26 G ½ hypodermic needle (Beckton Dickinson). Embryos were washed in 0.01% Tween 20 in PBS (PBS-T) and blocked with 10% BSA in PBS-T (blocking buffer) for 1 hour. Primary antibodies were diluted in a 50:50 mixture of blocking buffer:PBS-T (dilution buffer) and embryos were incubated for 2 hours at room temperature or overnight at 4 °C. To visualize RhoGEF2, we used embryos that expressed GFP-tagged RhoGEF2 under an endogenous promoter, which was recognized with an anti-GFP antibody (produced by our lab) diluted at 1:500. F-actin was visualized by incubating embryos with Alexa Fluor 647-conjugated phalloidin (Invitrogen) in dilution buffer. Secondary antibodies against the rabbit anti-GFP antibody was conjugated with Alexa Fluor 488 (Invitrogen) diluted at 1:500 in dilution buffer and incubated for 2 hours at room temperature or overnight at 4 °C. After incubations, embryos were mounted onto glass slides using AquaPolymount (Polysciences) and dried overnight.

All images were taken on a Zeiss LSM 710 confocal microscope with a 40x/1.2 Apochromat water objective lens, an argon ion, 561 nm diode, 594 nm HeNe, 633 HeNe lasers, and Zen software. Pinhole settings ranged from 1 – 2.5 airy units. For two-color live imaging, band-pass filters were set at ∼490 – 565 nm for GFP and ∼590 – 690 nm for mCH. For three-color imaging, band-pass filters were set at ∼480 – 560 nm for Alexa Fluor 488 and ∼660 – 750 nm for Alexa Fluor 647.

### dsRNA injections

To generate dsRNA that targets *trbl* transcripts for RNAi, the following primers were used to generate ∼200-base pair fragment: forward, 5’-TAA TAC GAC TCA CTA TAG GGT GCA GTA TGA ATC ACT GGA AGG −3’, and reverse, 5’-TAA TAC GAC TCA CTA TAG GGC CAC CAA CAT GGT GTA CAG G-3’. Each primer contains a T7 sequence at its 5’ end for use with the MEGAshortscript T7 transcription kit (Thermo Fisher Scientific). The reaction was placed in boiling water and allowed to cool to room temperature to promote annealing. RNA was extracted with phenol:chloroform, washed with ethanol, and resuspended in injection buffer (0.1x phosphate-buffered saline in DEPC water).

Dechorionated embryos were mounted onto glass slides and desiccated for 4 minutes using Drierite (Drierite). Embryos were covered with a 3:1 mixture of halocarbon 700/halocarbon 27 oils and then injected laterally with dsRNA in injection buffer into stage 2 embryos. As a control, injection buffer was injected. After injection, excess oil was wicked off and slides were prepared for live imaging. Embryos were incubated at 25 °C until they had completed cellularization.

### Image processing and analysis

All images were processed using MATLAB (MathWorks) and FIJI (http://fiji.sc/wiki/index.php/Fiji). A Gaussian smoothing filter (kernel = 1 pixel) was applied. Apical projections are maximum intensity Z-projections of multiple z sections (2-4 μm) and sub-apical sections are optical slices that are 1 – 2 μm below the apical sections.

Image segmentation for quantifications of cell area and anisotropy as well as myosin intensities was performed using custom MATLAB software titled EDGE (Embryo Development Geometry Explorer; https://github.com/mgelbart/embryo-development-geometry-explorer; Gelbart et al., 2012). Cell boundaries were automatically detected and manually corrected, after which EDGE exported cell area and anisotropy data. Cell apex anisotropy is calculated by fitting an ellipse to each cell. This measurement is calculated relative to the embryonic anteroposterior (AP) and dorsoventral (DV) axes. The length from the center of the ellipse to the edge along the AP axis is divided by the length from the center to the edge along the DV axis. For Figure 5D, we quantified cell aspect ratio, which is calculated from the geometry of the fitted ellipse. The aspect ratio is defined as the length of the major axis from the centroid to the edge of the ellipse over the length of the minor axis from the centroid to the edge. For the cell aspect ratios (Figure 5D) and cell area analysis of mitotic domain and non-mitotic domain cells (Figure 5E), we smoothed the data for each cell by a moving average (5 time steps wide). For the myosin intensity quantification in Figure 2F, medial myosin intensity was measured in EDGE as the total integrated pixel intensity of Myo::GFP signal at the apical cortex, excluding the segmented cell boundary.

To calculate the average medial myosin intensity before and during division in *arm* mutants (Figure 3D), the apical intensity of myosin in the cell was calculated with EDGE, as described above. The myosin intensities of mitotic cells were measured when the nuclear envelope had broken down. For these same cells, myosin intensity before division was measured 7 minutes prior to nuclear envelope breakdown. To calculate the ratio of apical:basal Baz::GFP intensities (Figure 3E), orthogonal (x-z) images were created for individual cells. A 2.5 μm by 17 μm region of interest was specified and the maximum pixel intensity within the region was calculated. This was done for both apical and basolateral regions, where the basolateral region was defined as being 17 μm lower than the apical region. The mean background fluorescence was subtracted from the maximum pixel intensities of the apical and basal regions for each cell. The ratio of apical to basal intensity was then calculated by dividing the corrected apical intensity by the corrected basal intensity. The ratio of average cytoplasmic to junctional RhoGEF2 intensity (Figure 4D) was measured as described above for apical:basal Baz::GFP intensities. First, orthogonal images were created. To acquire the intensity of RhoGEF2, a .5 μm by 7 μm region of interest in the apical region of the cell membrane was specified and the average pixel intensity within the region was measured. The cytoplasmic region was defined as the middle of the cell, excluding the nucleus, and average RhoGEF pixel intensity was measured in a 3.5 μm by 1 μm region. To calculate the ratio, the cytoplasmic value was divided by the junctional value.

## Acknowledgments

We would like to thank members of the Martin laboratory for their helpful comments and suggestions on the project. We would also like to thank Iain Cheeseman and Becky Lamason for their suggestions on the project and comments on a draft of this manuscript. Finally, we thank the Bloomington Stock Center and the TRiP at Harvard Medical School (NIH/NIGMS R01-GM084947) for providing fly stocks used in this study. This work was supported by National Institute of General Medical Sciences grant R01-GM125646 to A. C. Martin.

## Author contributions

C.S. Ko and A.C. Martin conceptualized the project and designed experiments. C.S. Ko and P. Kalakuntla performed the experiments. C.S. Ko and P. Kalakuntla analyzed the data. C.S. Ko and A.C. Martin wrote the manuscript. All authors reviewed and approved the final version of the manuscript.

---

**Table S1.**

| **Stock** | **Genotype** | **Source** |
| --- | --- | --- |
| 1 | w; P{w+ Ubi-H2A::GFP}3 | Bloomington Drosophila Stock Center |
| 2 | w; gap43::mCherry-7/TM3 | Lab stock |
| 3 | y, w; P{w+ sqh::GFP}42; Df(3L)ri-79c/TM3, Sb[1] | This study |
| 4 | y, w; P{w+ sqh::GFP}42; P{EP}trblEP3519/TM3, Sb[1] | This study |
| 5 | y, w; sqh::mCherry, Utr-ABD::GFP; P{EP}trblEP3519/TM3, Sb[1] | This study |
| 6 | UAS-fog 9A/TM3, Sb, Ser | Dawes-Hoang et al., 2005 |
| 7 | y, sc, v, sev; P{TRiP.GL01052}attp2 | DRSC/TRiP |
| 8 | w; mat67, Sqh::GFP; mat15, Gap43::mCherry/TM3, Sb[1] | Vasquez et al., 2014 |
| 9 | y, w; mat67, Sqh::mCherry; mat15, E-cadherin::GFP | Mason et al., 2016 |
| 10 | arm043A01, FRT101/FM7; sqh::GFP | Martin et al., 2010 |
| 11 | ovoD1, FRT101; hsFLP | Martin et al., 2010 |
| 12 | y, w; sqh>rok(K116A)::GFP; Df(3L)ri-79c/TM3, Sb[1] | Sqh>rok (K116A) is gift from J. Zallen, S. Simoes, and R. Fernandez-Gonzalez |
| 13 | y, w; sqh>rok(K116A)::GFP; P{EP}trblEP3519/TM3, Sb[1] | Sqh>rok (K116A) is gift from J. Zallen, S. Simoes, and R. Fernandez-Gonzalez |
| 14 | w; P{UAS-stg}/CyO | Bloomington Drosophila Stock Center |
| 15 | w; mat67, UAS-BazGFP[r1]/CyO | Eric Wieschaus (Wang et al., 2012) |
| 16 | w; P{sqh-pbl-EGFP}30 | Bloomington Drosophila Stock Center |
| 17 | y, w; Df(3L)ri-79c/TM3, Sb[1] | Bloomington Drosophila Stock Center |

| **Figure** | **Stocks used**<br>**(Female x Male)** |
| --- | --- |
| 1 | 1 x 2 |
| 2A | 5 x 17 |
| 2B | 3 x 4 |
| 2D-F; 4C-D;<br>5A-E | 6 x 8 |
| 3A-D | 10 x 11 |
| 3E-G | 15 Sibling Cross |
| 4A-B | 12 x 13 |
| 4E | 16 Sibling Cross |
| 5C-E | 7 x 9 |
| S1 | 14 x 8; 3 x 4 |
| S2 | 7 x 9 |
| S3 | 6 x 8; 7 x 9 |

## Supplementary Information

**Video 1. *Trbl* RNAi causes premature cell divisions in the mesoderm.** Embryos expressing Histone:GFP (H2A; green) and Gap43::mCH (magenta) injected with buffer (top) or dsRNA (bottom). Images were acquired every 15 seconds (top) or 22 seconds (bottom) and videos are displayed at 15 frames per second. Bars, 20 μm.

**Video 2. Apical myosin is lost during mitosis in *trbl* mutants.** Trans-heterozygous embryo (Df/EP3519), which displays the *trbl* phenotype, expressing Myo::GFP. Images were acquired every 6 seconds and video is displayed at 20 frames per second. Note that apical myosin is lost and then returns after mitotic exit. Bars, 20 μm.

**Video 3. Ectopic expression of *fog* in the ectoderm.** Embryos with ectopic *fog* overexpression expressing Myo::GFP (green) and Gap43::mCH (magenta). Medioapical myosin is lost in mitotic cells and ectopic furrows form between mitotic domains. Images were acquired every 17 seconds and video is displayed at 15 frames per second. Bars, 20 μm.

### Supplementary Figures

**Figure S1.**
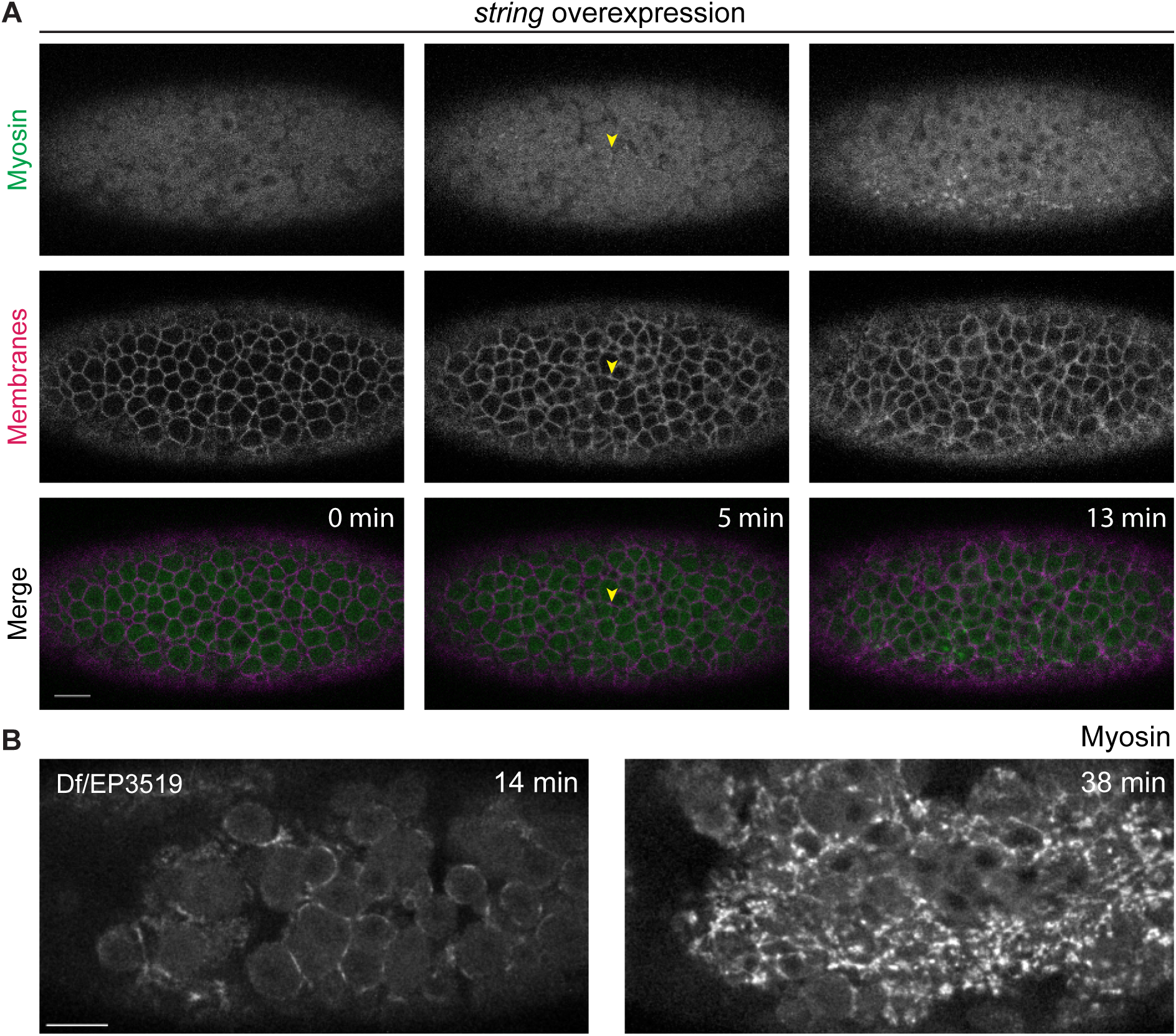
Overexpressing *string* (CDC25) results in the completion of cycle 14 divisions before ventral furrow formation. (A) Images are maximum intensity projections of a live *string* overexpressing embryo with Myo::GFP and Gap43::mCherry. A cytokinetic furrow from a premature division in the mesoderm is highlighted by a yellow arrowhead. The embryo is slightly rotated and ventral is towards the bottom. (B) Medioapical myosin returns in ventral cells of *trbl* embryos after completion of mitosis. Images are maximum intensity projections of a projections from a live trans-heterozygous embryo (Df/EP3519) expressing Myo::GFP. Scale bars, 15 μm.

**Figure S2.**
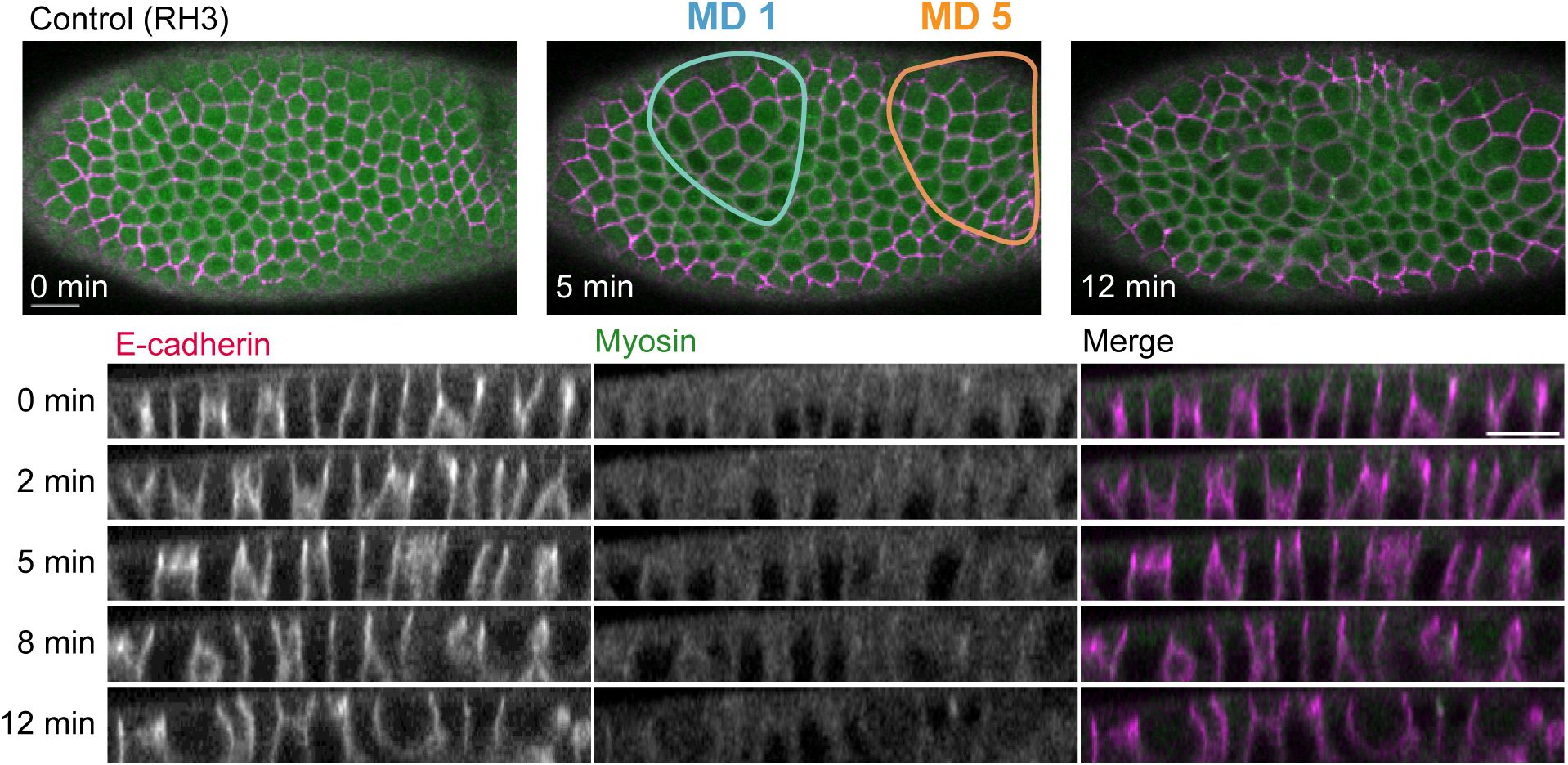
Furrows do not normally form between mitotic domains and mitotic cells do not normally elongate in the dorsal head. Images are maximum intensity projections from a live control embryo (Rhodopsin 3 shRNA line) expressing Myo::CH and E-cadherin::GFP. Image acquisition field is slightly more anterior; the cephalic furrow lies posterior to mitotic domain 5. Scale bars, 15 μm.

**Figure S3.**
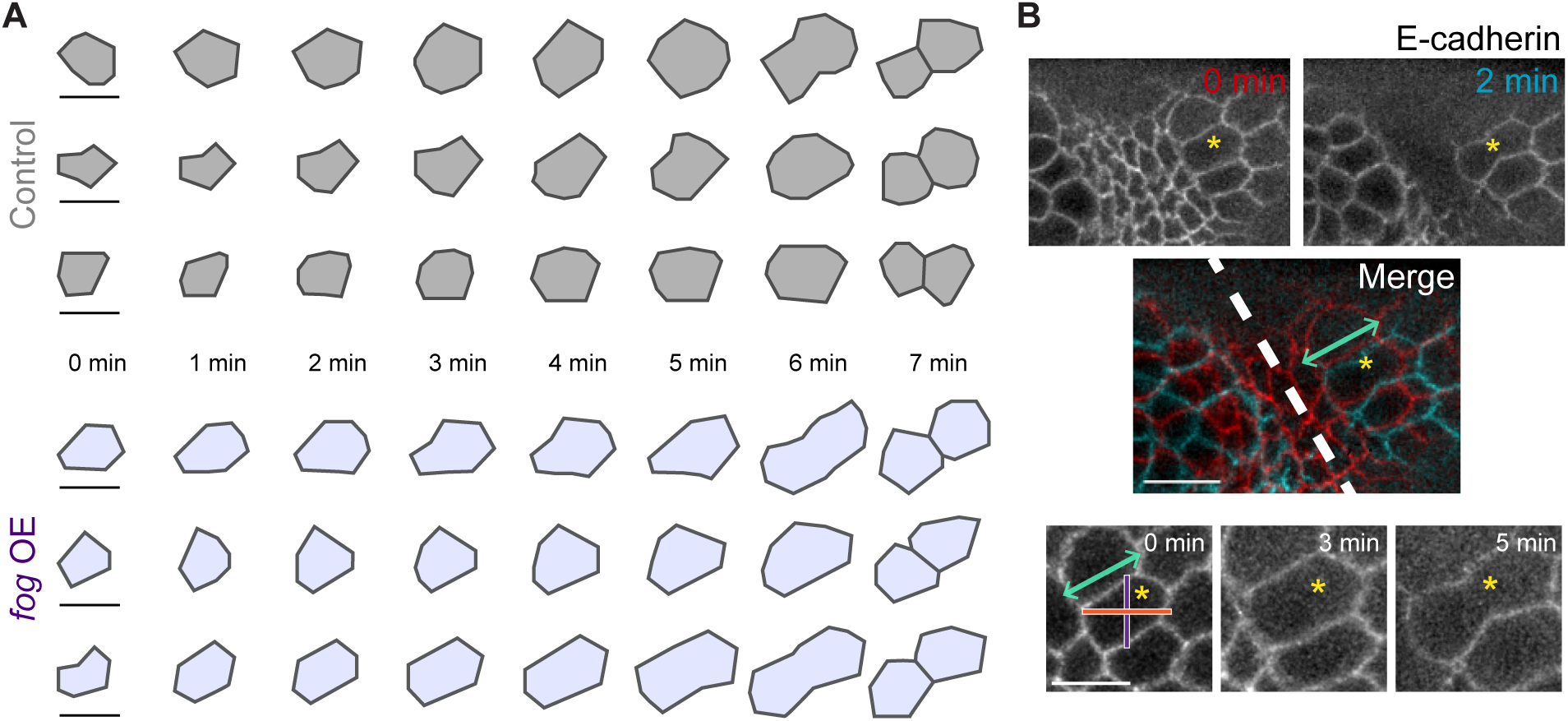
Mitotic cells are more elongated when *fog* is ectopically expressed. (A) Representative mitotic domain 3 cells from control (Rhodopsin 3 shRNA line) and ectopic *fog* expression embryos are segmented and shown here. Initial cell size was set 7 minutes prior to cytokinetic ring formation. (B) Mitotic cells become stretched towards the ectopic furrow (dashed line). Images are maximum intensity projections of *fog* overexpressing embryos expressing E-cadherin::GFP. The same cell is highlighted by the yellow asterisk. The axis of stretch is indicated by the double-sided arrow. The body axes of the embryo (AP, orange; DV, purple) are shown on the bottom set of images. Scale bars, 15 μm (B, top), 10 μm (A and B, bottom).

